# Memory reactivation during sleep does not act holistically on object memory

**DOI:** 10.1101/2023.12.14.571683

**Authors:** E.M. Siefert, S. Uppuluri, J. Mu, M.C. Tandoc, J.W. Antony, A.C. Schapiro

**Affiliations:** Department of Psychology, University of Pennsylvania, Philadelphia, PA, 19104, USA; Department of Neurosurgery, Perelman School of Medicine, University of Pennsylvania, Philadelphia, PA, 19104, USA; Department of Psychology and Child Development, California Polytechnic State University, San Luis Obispo, CA, 93407, USA

## Abstract

Memory reactivation during sleep is thought to facilitate memory consolidation. Most sleep reactivation research has examined how reactivation of specific facts, objects, and associations benefits their overall retention. However, our memories are not unitary, and not all features of a memory persist in tandem over time. Instead, our memories are transformed, with some features strengthened and others weakened. Does sleep reactivation drive memory transformation? We leveraged the Targeted Memory Reactivation technique in an object category learning paradigm to examine this question. Participants (20 female, 14 male) learned three categories of novel objects, where each object had unique, distinguishing features as well as features shared with other members of its category. We used a real-time EEG protocol to cue the reactivation of these objects during sleep at moments optimized to generate reactivation events. We found that reactivation improved memory for distinguishing features while worsening memory for shared features, suggesting a differentiation process. The results indicate that sleep reactivation does not act holistically on object memories, instead supporting a transformation process where some features are enhanced over others.

## INTRODUCTION

Memory consolidation is the process of converting newly encoded information into long-term memories, and sleep is thought to be a critical time for this consolidation. In rodents, neural firing patterns during sleep recapitulate those observed during previous awake learning (Pavlides and Winson, 1989; Wilson and McNaughton, 1994), with disruptions to this reactivation impairing subsequent behavior (Girardeau et al., 2009; Ego-Stengel and Wilson, 2010; Klinzing et al., 2019; Aleman-Zapata et al., 2022). Research in humans has likewise illustrated that the repeated reactivation of memories for specific facts, objects, and associations during sleep can improve memory for these items (Oudiette and Paller, 2013; Hu et al., 2020; Schreiner and Staudigl, 2020).

This simple picture suggests that memories are reactivated in whole during sleep and that the entirety of the memory benefits from this reactivation. But we know that memories are not holistically stabilized and strengthened across time. Rather, they are transformed, with individual features of memories differentially strengthened and weakened (Schacter et al., 2011; Winocur and Moscovitch, 2011; Brady et al., 2013). For example, a week after visiting an aquarium, you may remember that a puffer fish was round and had spikes but forget what its tail looked like. Brady et al. (2013) demonstrated that different features of objects (color, state of being open/closed) are forgotten independently over time, arguing that memory changes are not holistic because multi-featural memories are not always bound into individual units (c.f., Balaban et al., 2020). Additionally, a large body of work has demonstrated that unique details of an encoded event can be forgotten while memory for the general structure of the event is retained (Winocur and Moscovitch, 2011). Sekeres et al. (2016, 2018b) showed that a week after encoding detail-rich videos, participants remember only the central, gist elements of some videos while retaining the peripheral details of other videos. The mechanisms driving these selective changes to memory features remain poorly understood. In this work, we asked whether sleep-dependent reactivation plays a role in memory transformation over time.

The Complementary Learning Systems (CLS) framework proposed that interleaved reactivation of memories offline, perhaps especially during sleep, supports a transformation in the nature of the memories (McClelland et al., 1995). According to this framework, memories are initially stored in an orthogonalized manner in the hippocampus and then interleaved offline replay supports the extraction of the shared structure across these memories in neocortical areas. The increased reliance on neocortical areas, employing more overlapping representations, supports a transformation of these memories that emphasizes their shared over their unique features. The Trace Transformation Theory posits a similar transformation in memory quality accompanying a change in underlying memory system (Winocur and Moscovitch, 2011; Sekeres et al., 2018a). Whether reactivation during sleep supports these kinds of memory transformations, and whether the interleaved order of reactivation in particular is important (as posited by CLS), remain unanswered questions.

To address how reactivation may transform structured memories (i.e., memories containing multiple features with structured interrelationships), we applied Targeted Memory Reactivation (TMR) after participants learned categories of novel objects. TMR allows causal manipulation of sleep reactivation in humans, where different sounds (Rudoy et al., 2009) or odors (Rasch et al., 2007) are associated with information during learning and then re-presented during sleep to cue the reactivation of the associated information (Oudiette and Paller, 2013; Lewis and Bendor, 2019; Hu et al., 2020). Administering TMR cues during sleep has been demonstrated to increase replay of associated memories in rodents (Bendor and Wilson, 2012) and reactivation of task-specific content in humans, with benefits to memory (Belal et al., 2018; Cairney et al., 2018; Wang et al., 2019; Abdellahi et al., 2023).

Here, we administered TMR to examine whether reactivation impacts object memory holistically or whether it supports memory transformation, with some features benefitting more than others. Participants learned categories of novel objects that each had multiple visual features and an auditorily-presented name. We then used these names as auditory TMR cues during a subsequent nap. We administered the cues during NREM sleep in different orders across object categories (interleaved vs. blocked) to assess whether the order of reactivation impacts later memory. We found that reactivation during sleep improved memory for distinguishing item features while worsening memory for features shared across members of a category. The improvement to distinguishing features was especially strong for items reactivated in blocked order, and the benefit transferred to uncued items from the same category as cued items. Our results provide evidence that reactivation during sleep does not act holistically on structured object memories and instead can enhance certain features of memories at the expense of others.

## MATERIALS AND METHODS

### Participants

Thirty-four participants (mean age (SD): 20.79 (2.45); range: 18-29; 20F) recruited from the University of Pennsylvania community took part in the experiment. All participants reported no history of neurological or sleep disorders and normal sleep schedules (overnight sleep of at least six hours with bedtimes between 10pm and 2am). All participants had normal or corrected-to-normal vision, normal hearing, and were fluent in English. To prepare for the study, participants abstained from alcohol, recreational drugs, psychologically active medications, and caffeine from 24 hours prior. Data from nineteen additional participants were excluded due to: inability to reach 50% accuracy during learning (*N* = 2), no sound cues being administered (did not reach N2 or N3 sleep; *N* = 9), reported hearing sounds in post-nap survey (*N* = 4), and other technical problems (*N* = 4). The study was approved by the local Institutional Review Board and informed consent was obtained from all participants. Participants received either monetary payment or course credit as compensation.

### Stimuli

Stimuli were adapted from Schapiro et al. (2017): Participants learned about fifteen novel “satellite” objects that were organized into three categories (alpha, beta, and gamma). Each satellite possessed an individual name and five visual parts. One of the satellites in each category contained all prototypical parts for that category. All other satellites contained four prototypical parts and one part that was completely unique. Thus, satellites had shared features (prototypical parts) as well as unique features (Fig. 1A). Satellites were constructed randomly for each participant, following this category structure. Across the experiment, satellite objects were presented in black and white on a white background, subtending a visual angle of approximately 19 degrees.

**Figure 1.**
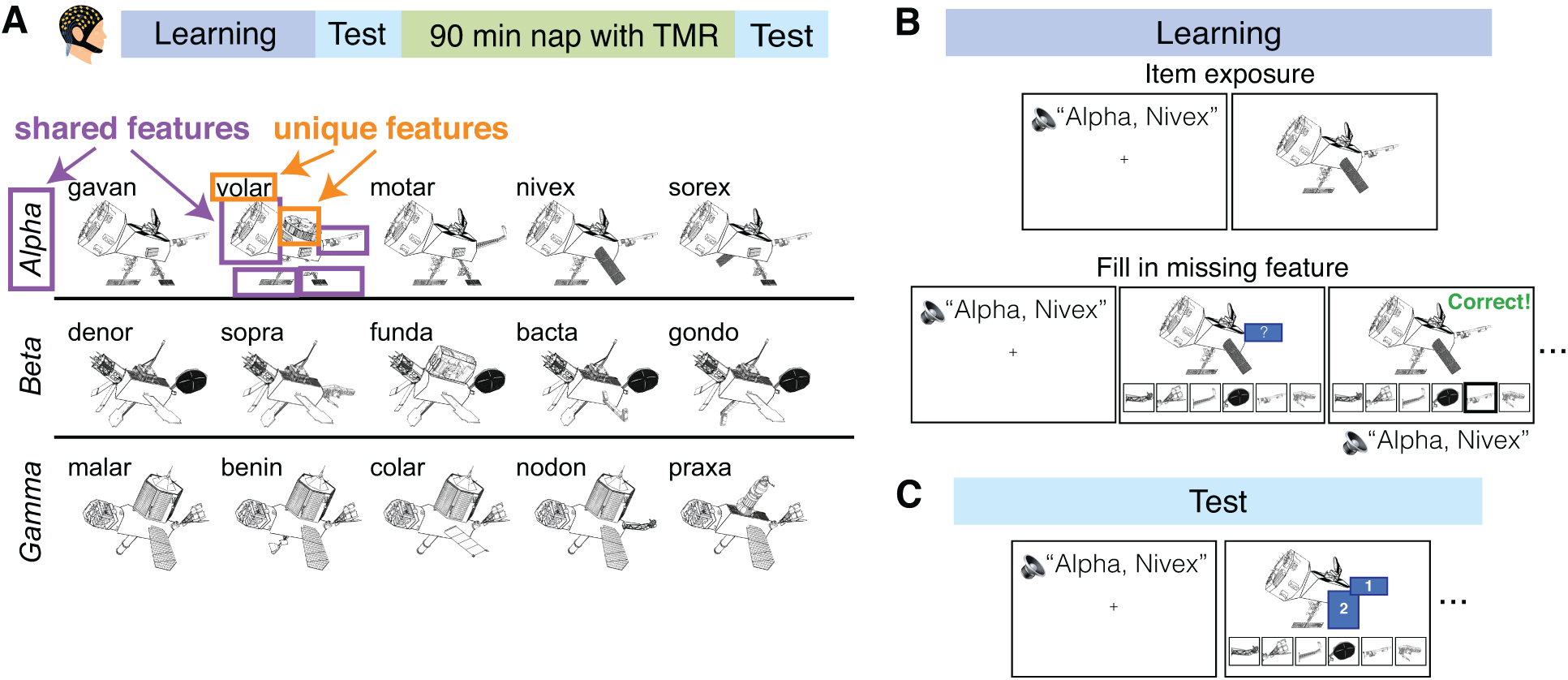
Experimental Design. ***A*,** Study timeline and stimuli. Participants completed a novel object category learning task and took a nap where Targeted Memory Reactivation (TMR) was performed. They studied three categories of novel “satellite” objects (alpha, beta, gamma), where a satellite could have parts that were unique to itself and parts shared with other members of its category. Each satellite had its own unique name (ex: nivex). ***B*,** Learning phase. First, participants were exposed to the satellites one-by-one: participants heard a satellite’s name out loud and then saw it on screen. Next, participants completed blocks of trials where they first heard a satellite’s name and then were shown a satellite with one feature missing and instructed to select the missing feature. ***C*,** Test phase. Participants heard a satellite’s name and then were shown a satellite with one or two features missing. They selected a feature and then rated their confidence in their decision.

**Figure 2.**
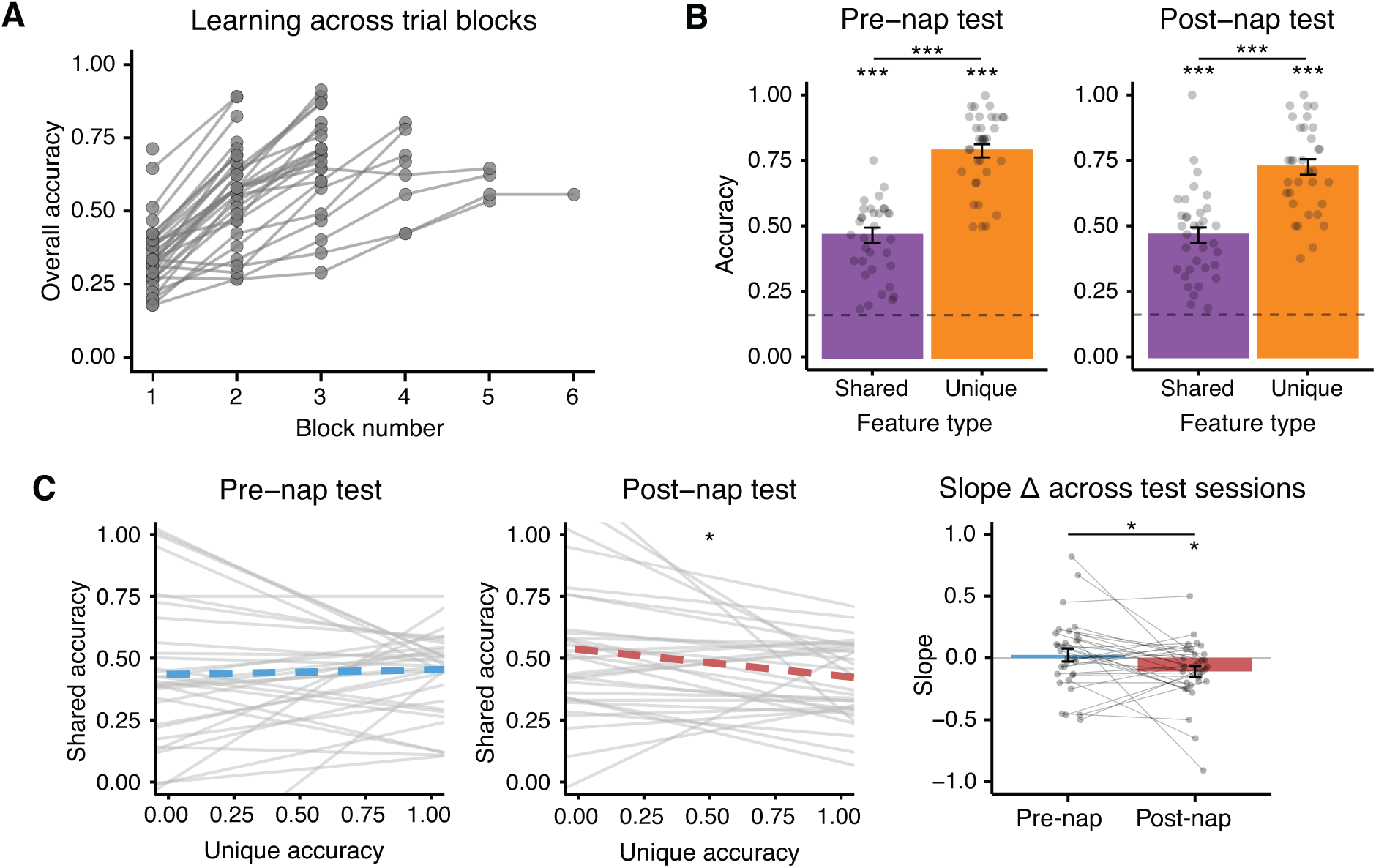
Pre- and post-nap behavior. ***A*,** Overall accuracy across learning blocks. ***B*,** Pre-nap and post-nap test performance. Mean accuracy on unique (orange) and shared (purple) feature memory trials. The dotted line indicates chance. ***C*,** Assessment of tradeoff in unique and shared feature accuracy. A line was fit, for each participant, predicting shared feature accuracy from unique feature accuracy of the corresponding object, for the pre-nap (left) and post-nap (middle) tests. Mean slope across participants is represented with a thick dotted line. A negative slope indicates a tradeoff in shared and unique feature accuracy. Right: Barplot of slopes in the pre-nap (blue) and post-nap (red) tests. Bars represent mean slopes across participants. Error bars represent +/- 1 SEM. Dots and lines correspond to individual participants. \**p* < .05, \*\**p* < .01, \*\*\**p* < .001

### Experimental Design

The experiment consisted of one five-hour session which was split into four phases: learning, pre-nap test, nap period with Targeted Memory Reactivation (TMR), and post-nap test (Fig. 1A). The learning and test phases of this experiment were adapted from Schapiro et al. (2017).

Participants arrived at the laboratory between 12:30 and 1:00pm. During consent, they were informed that they would be taking part in a memory and nap study, but were not informed that TMR would occur. Next, participants completed the learning task and pre-nap test, and then were given a 90 min nap opportunity in the lab, when TMR was administered. After waking up, participants were given 20 mins to complete a post-nap survey and overcome sleep inertia. Participants were excluded from analyses if they reported hearing satellite sounds during the nap on this survey. Next, they completed the post-nap test. Finally, participants completed an exit survey regarding their learning and test strategies. Behavioral tasks were administered with MATLAB 2019a using the Psychophysics Toolbox (Brainard, 1997; Pelli, 1997; Kleiner et al., 2007), and surveys were administered through REDCap.

#### Learning task

First, the fifteen satellites were introduced to the participant. For each satellite, an audio recording of its category name and individual name was played. Then, the satellite’s image was displayed for eight seconds (Fig. 1B).

Next, participants completed learning trials where they again heard a satellite’s category and individual name and then viewed an image of the satellite. However, this time the satellite images had one visual feature covered up. Participants chose from six feature options to complete the satellite (Fig. 1B). If the participant chose the correct feature, they were told they chose correctly and presented with the satellite image and audio name again. If the participant chose the incorrect feature, they were shown the correct feature and repeated the trial until choosing correctly. Unique features were queried 2.21 times more frequently than shared features, based on pilot testing aimed at matching performance across shared and unique features. (Though our aim was to match performance, and pilot results suggested matched performance at this ratio, our final sample had much better memory for unique features.)

Learning trials were organized into blocks of 45 trials. Participants continued the task until achieving 66% correct in a block or 50% in a block after 60 min of learning (six participants reached 50% but not 66% in 60 min of learning).

#### Pre-nap and post-nap test

Participants were tested on shared and unique features of the satellites as well as on novel satellites (satellites that belonged to the studied categories but had one completely novel feature). Participants were told that a select number of satellites could be new and to complete them to the best of their ability.

The test was very similar to the learning phase but contained no feedback: After hearing a satellite’s category and individual name, participants were shown the satellite on screen with one or two features covered up with numbered boxes. They then filled in each box using six available options (Fig. 1C). Participants were tested twice on each prototypical satellite, where each time they were queried on two different shared features. For all other studied satellites, participants were tested on (a) two shared features, (b) a shared feature and then the unique feature, and (c) the unique feature and then a shared feature. Across these three test trials, the unique feature of a satellite was queried twice and the shared features were each queried once. When a novel satellite was tested, the feature queried was always one shared feature. The satellite order was pseudorandomized across trials such that no satellite was tested twice in a row. After choosing a feature, participants rated their confidence level on a scale of one to five (one being very unsure; five being very sure). The confidence question was added after the first eight participants (thus, 26 of 34 participants completed confidence judgments). The pre-nap and post-nap tests were identical but with shuffled trial orders.

#### Nap period with TMR

Participants were provided a 90-min opportunity to nap beginning at ∼2:00-2:30 PM. All lights and monitors were turned off and white noise was played through speakers in the nap room to help mask any external sounds (40 dB, measured at pillow).

Once a participant reached N2 sleep, the individual satellite names were played using our real-time protocol to encourage the reactivation of associated satellite information (Targeted Memory Reactivation, TMR). Sounds were played over the white noise and resulted in increases of no larger than 3 dB (measured at pillow).

We administered the TMR sound cues in a different order for each category of satellites: exemplars from one category of satellites were cued in interleaved order, exemplars from a second category in blocked order, and one category was left uncued (Fig. 3A). Satellite categories were randomly assigned to cueing orders, and there were no significant pre-nap accuracy differences for items assigned different cueing orders for shared (one-way ANOVA: f(2) = .75, p = .48) or unique (f(2) = .19, p = .31) features. Additionally, one randomly chosen exemplar from each cued category was left uncued (uncued items in cued categories). The presentation of the two cued categories was intermixed across N2 and N3 sleep, and cueing was stopped immediately upon indication of N1, wake, or REM. Cues were resumed if the participant re-entered N2 or N3. Items were cued between one and eighteen times (mean = 7.59, SD = 2.90) depending on how much time the participant spent in N2/N3 sleep (Extended Data Figure 3-1). Throughout the nap, the researcher sat in an adjacent control room with audio and EEG monitoring and an intercom system to communicate with the participant.

**Figure 3.**
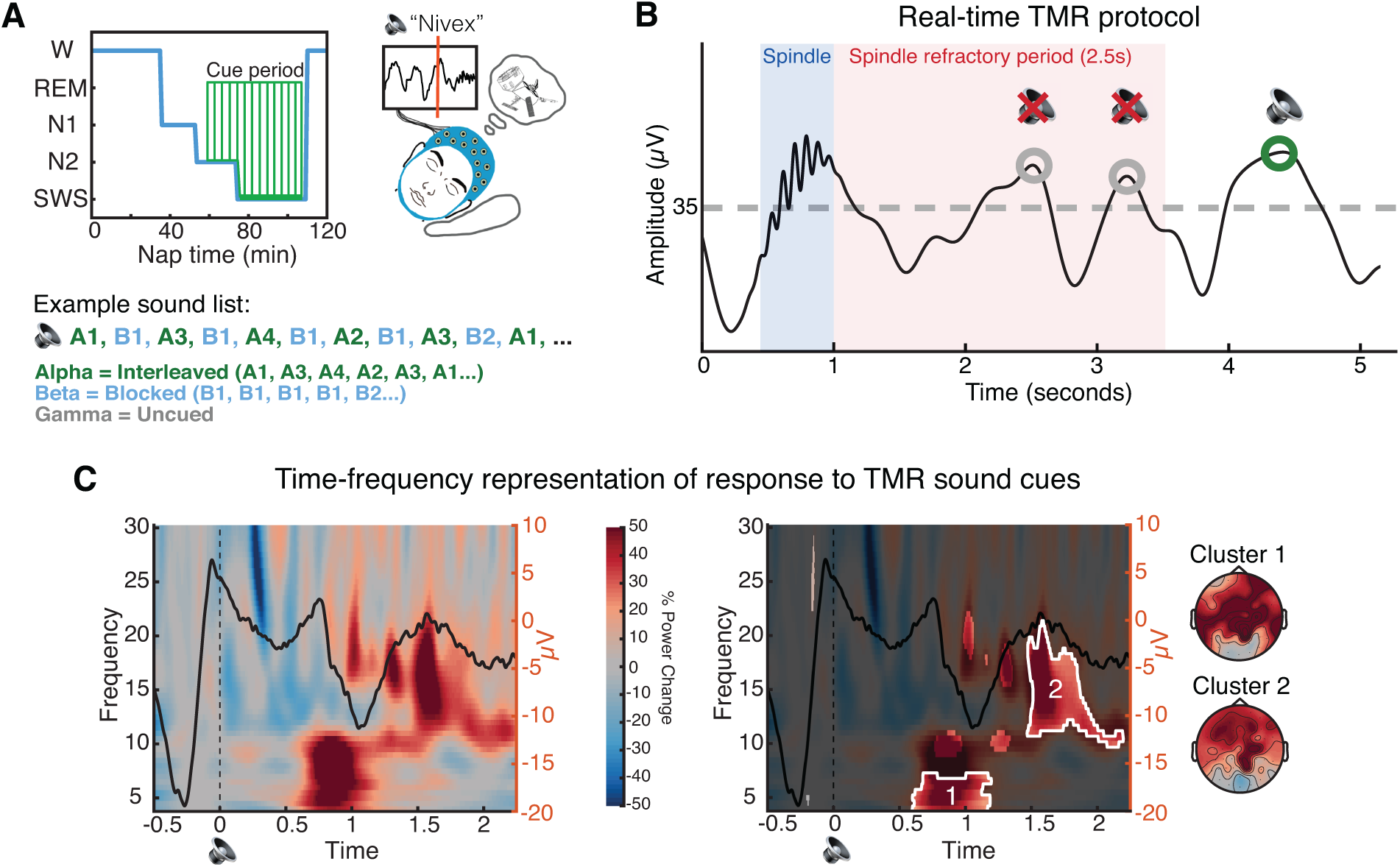
Real-time Targeted Memory Reactivation protocol. ***A*,** Overview of Targeted Memory Reactivation (TMR). (left) TMR cues — delivered aloud as individual satellite names — began after a participant entered and remained in N2 sleep for at least three minutes. One category was cued in interleaved order, one in blocked order, and one was left uncued. The presentation of the two cued categories was intermixed across NREM sleep. See Extended Data Figure 3-1 for more details. ***B*,** Real-time TMR administration. We developed a novel TMR protocol in which cues were played in the up-states of slow oscillations (up-state = signal goes above threshold of +35 μV; Ngo and Staresina, 2022) and at least 2.5 seconds after a detected spindle (Antony et al., 2018b). ***C*,** left: Time-frequency representation of the difference between the neural response when a TMR sound cue was played versus not (sham), with the ERP response to sound cues overlaid (averaged across all channels and all participants). Sounds were played at time = 0. Right: Cluster-based permutation testing identified two significant clusters in the time-frequency response to TMR sound cues. Un-shaded areas represent clusters identified via performing *t*- tests across participants (α = .01). Clusters that survived subsequent permutation testing are highlighted in white. Topoplots show the spatial representation of the identified clusters. See Extended Data Figure 3-2 for more details.

### Sleep EEG recording

EEG was recorded during learning, test, and nap phases via an actiCHamp amplifier (Brain Vision) using a 64-channel EasyCap EEG cap. Data were obtained with a sampling rate of 512 Hz and an online reference placed at FCz. For the purposes of sleep-scoring, two electrodes were repurposed to record electrooculography (EOG), two to record electromyography (EMG), and two were placed on the mastoid. EEG capping took place at the start of the experiment (before the learning phase), and impedance was checked at the start as well as right before and after the nap (impedance of all electrodes was kept below 20 kΩ).

### Real-time targeted memory reactivation

A growing body of literature has suggested that endogenous reactivation occurs in slow-oscillation up-states (Diekelmann and Born, 2010; Mölle and Born, 2011; Gulati et al., 2014; Lewis and Bendor, 2019; Schreiner et al., 2021), and administering Targeted Memory Reactivation (TMR) cues in slow-oscillation up-states reliably leads to memory improvements (Batterink et al., 2016; Göldi et al., 2019; Ngo and Staresina, 2022; Abdellahi et al., 2023; Xia et al., 2023). Additionally, reactivation may be mediated by spindles, which are often nested within these up-states (Born and Wilhelm, 2012; Klinzing et al., 2019) and which exhibit refractory periods (Antony et al., 2018b). TMR cues administered immediately after detected spindles have been shown to disrupt memory for cued items, perhaps because they occurred in this post-spindle refractory period (Antony et al., 2018b). Here, we developed a real-time protocol to play TMR cues both in the up-states of slow-oscillations and outside of the spindle refractory period (2.5s).

Real-time TMR phase-locked to slow-oscillation (SO) up-states and with reference to spindle timing was performed using the OpenViBE Brain Computer Interface Software (Renard et al., 2010) and custom MATLAB scripts. The protocol we developed was a combination of two previously published systems: SO up-state detection as described in Ngo & Staresina (2022) and online spindle detection as described in Antony et al. (2018b). The TMR system was turned on by an experimenter after a participant was in N2 sleep for at least three minutes. Once turned on, the system ran the two processes (SO, spindle detection) in parallel, with the time of spindle occurrence being sent from the spindle detection to the SO detection. Specifically, once a SO was detected (Fz filtered signal rising to cross a threshold of +35 μV), the system monitored how much time had passed since the time of the last spindle. If it had been at least 2.5 seconds since the last detected spindle, the system played a sound once a SO upstate was detected (signal changing slope from positive to negative, indicating a local maximum). Otherwise, the system returned to looking for a SO until 2.5 seconds since the last spindle had passed (Fig. 3B). After playing a sound, the TMR detection system paused for eight seconds (median empirical time interval between cues: 9.56s, range: 8.04 - 3178.5s, with values at the high end due to temporary sleep state transitions out of N2/N3). In addition to playing sounds, spaced throughout the cueing list were ‘no sound’ triggers, in which the system did not play a sound but sent a trigger marking when it would normally play a sound. These ‘no sound’ marked SOs were then used as a sham for comparing the impacts of cueing to endogenous SOs. This system was implemented using MATLAB 2011a (32bit), OpenViBE Designer v2.2.0 (32bit), and OpenViBE for Brain Products server (32bit).

### Statistical Analysis

#### Behavioral data analyses

All analyses of behavioral data were performed in R (R Core Team, 2021). We examined the impacts of TMR cueing on memory for unique and shared satellite features using linear mixed effects models (*lme4* package; Bates et al., 2015). The main model applied to our data was:

> *model <- lmer(accuracy difference ∼ cueing * feature type + # times cued + pre-nap test accuracy + (1|subject), data)*

*Accuracy difference* was the numeric value of the accuracy change for an item and feature type (shared, unique) combination across the nap, resulting in two data points for each satellite for each subject (accuracy change for shared features, accuracy change for unique features). *Cueing* was a categorical variable with levels “cued” and “uncued”. Items were counted as uncued if they were in the pre-designated uncued category. Items were counted as cued if they were in the pre-designated cued category and cued at least once. *Feature type* was coded as a categorical variable with levels “shared” and “unique”. *# times cued* was a numeric, mean centered variable that corresponded to the number of times an individual item was cued. *Pre-nap test accuracy* was the numeric value of pre-nap accuracy for an item and feature type (shared, unique) combination. Subjects were included as random effects (intercepts). See Extended Data Figure 4-1 for details on the number of data points, items, and participants included in this model (and all other models).

Impacts of cueing on novel items were assessed using the same model but with only novel items included (*feature type* term was removed). For the novel items, *cueing* was determined by which category the novel items were part of (i.e., if the novel item was in a category that was cued, it was counted as cued) and # *times cued* reflected the number of times that items from the novel item’s category were cued.

The model was also applied to the subset of items where shared feature accuracy was greater than or equal to unique feature accuracy in the pre-nap test (Fig. 4B). Two participants had no items that were cued more than once where shared feature accuracy was greater than or equal to unique feature accuracy (*N* = 32 for this subset).

**Figure 4.**
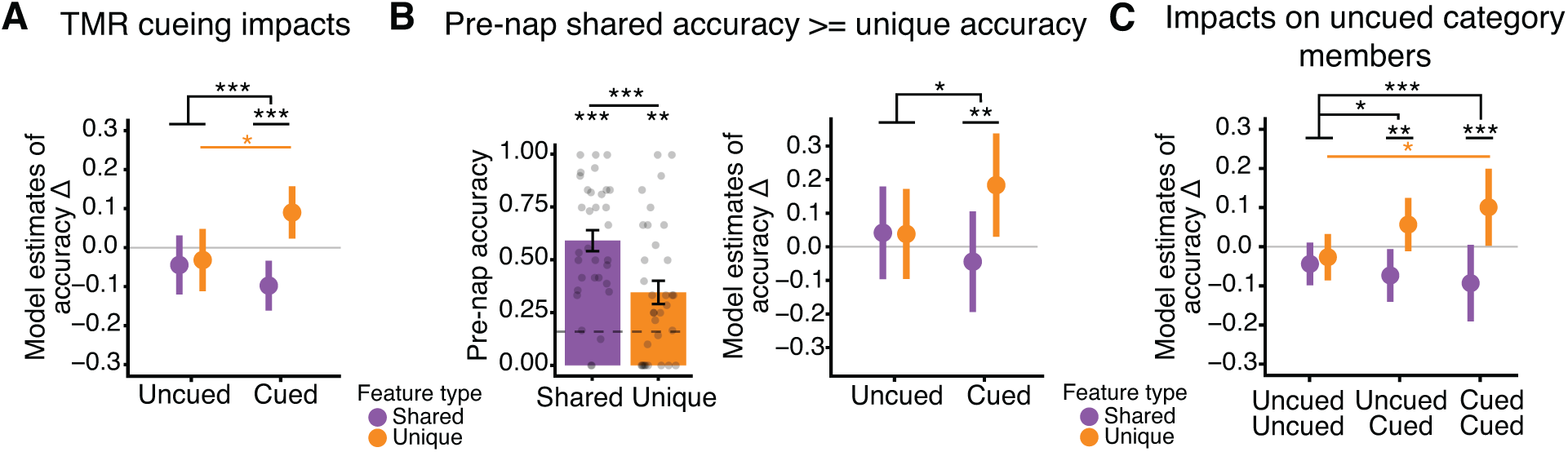
TMR cueing improved unique feature memory and impaired shared feature memory. ***A*,** Impacts of TMR. A linear mixed effects model was used to analyze change in unique and shared feature accuracy across the nap as a function of cueing. Model estimates for accuracy change for shared (purple) and unique (orange) features are plotted. ***B*,** Replication of analysis from A using only the subset of items whose shared feature accuracy was greater than or equal to their unique feature accuracy. Left: Mean shared and unique feature accuracy from the pre-nap test. Dots represent participants, error bars represent +/- 1 SEM, the dotted grey line represents chance. Right: Model estimates of unique and shared feature accuracy change for uncued and cued items from this subset. ***C*,** Impact of cueing on uncued items from cued categories. Plotted are the model estimates for unique and shared feature accuracy change for items in the designated “uncued” category (Uncued Uncued), items uncued in a designated “cued” category (Uncued Cued), and for items who were cued (Cued Cued). \**p* < .05, \*\**p* < .01, \*\*\**p* < .001. See Extended Data Figure 4-1 for additional model details.

To examine whether TMR cueing impacted items that were themselves uncued but in the same category as other cued items, we modified the *cueing* term of the model to include three levels: uncued items from uncued categories, uncued items from cued categories, cued items from cued categories (if cued at least once). We forced the exclusion of at least one item from cueing after the first few participants. Thus, two participants had no uncued items from cued categories.

To analyze how sleep stage impacted cueing-related memory changes, we modified the model’s *cueing* term to include four levels: items from uncued categories; items cued during only N2 sleep; items cued during only N3 sleep; items cued in both N2 and N3 sleep. Twenty participants had items cued in only N2 sleep; twenty-one had items cued in only N3 sleep; twenty-two had items that were cued in both N2 and N3 sleep; thirty-four had uncued items.

To assess the relationship between power in the identified theta and spindle band clusters and memory changes, we extracted, for each participant and item, the average power in each cluster (theta and spindle; see *Sleep EEG data analyses* for details). We then ran the model with the *cueing* term replaced by a numeric term representing either the power in the theta or spindle cluster. These models were run only on items that were cued.

To examine the effects of cueing order on memory, we modified the model’s *cueing* term to include two levels: interleaved and blocked. Since cueing an item just once would not allow for a proper distinction of blocked versus interleaved, items were included in the interleaved condition if they were from the designated interleaved category and cued four or more times; items were included in the blocked condition if they were from the designated blocked category and cued four or more times. In this model, we excluded uncued items to directly test for differences across blocked and interleaved cueing orders. We ran a second version of this model with uncued items included to confirm differences between the uncued condition and both the blocked and interleaved conditions. One participant was removed from this analysis due to a cueing order error and three additional participants did not have any items that were cued four or more times (*N* = 30).

To assess whether the order of items in blocked cueing had an impact on memory, we replaced the model’s *cueing* term with one of *sequence position*. This term ordered items by their relationship to the end of the nap, grouping them as follows: items from uncued categories (*sequence position* = 0), items cued fourth from last (1), items cued third from last (2), items cued second to last (3), items cued last (4). We treated this term as numeric to assess the overall trend across sequence positions, and then converted it to categorical to assess relationships between the individual positions. Only fully cued items from the blocked category were included in the cued groups in this analysis (we thus removed the *# times cued* variable from the model). We included items from uncued categories in these models as a baseline to compare cued items against. Twenty-seven participants had at least one item that was blocked and fully cued (sequence position 4); twenty-two participants had at least two items that were blocked and fully cued (sequence position 3); twenty-one participants had at least three items that were blocked and fully cued (sequence position 2); sixteen participants had at least four items that were blocked and fully cued (sequence position 1); twenty-seven participants had uncued items (sequence position 0).

For all models, we used the *Anova* function from the *car* package (Fox and Weisberg, 2019; Type II Wald chisquare tests) to assess the significance of model variables (provides significance of categorical variables). Then, the *lmerTest* package (Kuznetsova et al., 2017) was used to obtain p-values for models using Satterthwaite’s degrees of freedom method (provides significance for each level of categorical variables). The *emmeans* package (Lenth, 2022) was used to obtain estimated marginal means for post-hoc comparisons with Tukey’s adjusted p-values and degrees of freedom calculated using the Kenward-Roger method. Model assumptions of normality, linearity, and homoscedasticity were confirmed using *resid_panel* function from *ggResidpanel* package (Goode and Rey, 2022). Confirmation of effects using analyses of confidence data, rather than accuracy data, were performed using all of the same models and tests as applied to the accuracy data.

#### Sleep EEG data analyses

Analyses of sleep EEG data were performed in MATLAB (2022b) using the Fieldtrip toolbox (Oostenveld et al., 2011).

### Sleep stages

For sleep scoring, data were down sampled to 256 hz, re-referenced to the average of the linked mastoids, and filtered between .1 and 35 hz. Sleep scoring was done using Dananalyzer (https://github.com/ddenis73/danalyzer; Denis et al., 2021) by an expert scorer according to standard criteria (Iber et al., 2007).

### Time-frequency analyses

For both time-frequency representations (TFRs) and event-related potentials (ERPs), data were cut from -3 to 3 s centered on the time of cue delivery. Data were then down sampled to 256 hz, re-referenced to the average of the linked mastoids, and filtered between .3 and 30 hz. Artifacts were rejected in two stages: first, the experimenter manually looked through every individual trial and channel and made note of those which looked problematic. Next, trials and channels were confirmed for removal via FieldTrip’s visual summary functions. Trials and channels that were both noted as problematic during manual screening and also contained amplitude, variance, or kurtosis outliers were removed.

For TFR analyses, data were convolved with a 5-cycles Hanning taper and spectral power was obtained from 4-30 hz in .5 hz frequency steps and 5 ms time steps (Cairney et al., 2018). Then, participant-specific TFRs were converted into percent power change relative to a -300 ms to -100 ms pre-cue window (Cairney et al., 2018). For analysis of sound-elicited activity, TFRs for sound and no sound (sham) trials were processed separately and then a difference TFR was obtained for each participant (TFR sound minus TFR no sound; Fig. 3C). For the visualization of ERPs in Fig. 3C, data were high-pass filtered at .5 hz and baseline-corrected with respect to the 200 ms to 0 ms window before cue onset (Cairney et al., 2018).

To identify significant time-frequency clusters of sound-elicited activity, trial-specific spectrograms were averaged across electrodes within participants. We then performed a t-test across subjects, generating candidate clusters of points in time-frequency space where power was different from zero (α = 0.01). To account for multiple comparisons, we performed cluster-size based permutation testing: a permutation-corrected *p*-value was determined for each candidate cluster by calculating the frequency of the candidate cluster size exceeding maximum cluster sizes in a permutation distribution obtained by running one-sample t-tests after randomizing the sign of power values among participants (1000 permutations; significance level: *p* = .05).

In addition to analyzing time-frequency clusters of sound-elicited activity, we examined differences in sound-elicited activity for interleaved and blocked cues (interleaved TFR minus blocked TFR arrays), differences in elicited activity in N2 and N3 sleep (N2 TFR minus N3 TFR arrays; eight participants had no cues delivered in N3, *N* = 26) and differences in elicited activity across blocked sequence positions (last item cued TFR minus first item cued TFR; *N* = 22). We additionally used all of the same preprocessing and analysis steps for examination of any extended sound-elicited activity, cutting the data from -2 to 6 s centered on cue delivery.

## RESULTS

Participants came into the lab and learned three categories of novel satellite objects before taking a nap (Schapiro et al., 2017; Fig. 1A). Each satellite had its own unique name as well as visual features that could be unique to itself or shared with other members of its category. Participants completed learning trials where they heard a satellite’s name spoken and then were prompted to fill in missing features of the satellite, with feedback provided (Fig. 1B). After learning, participants were tested on their knowledge of the satellites in a similar feature inference test (Fig. 1C). Participants were tested on unique and shared features of the satellites, as well as on novel satellites from the studied categories. Participants then took a 90-minute nap in the lab during which the names of the satellites were played as cues for Targeted Memory Reactivation (TMR). Participants repeated the test after the nap.

### Pre-nap training led to especially strong memory for unique features

Participants trained for an average of 3.06 (SD = 1.07) blocks of learning trials (45 trials in a block) with the average proportion correct on the last block being .73 (SD = .10; Fig. 2A). In both the pre-nap and post-nap tests, participants were well above chance (0.167) on their memory for both unique (pre-nap: mean = .79, *t*[33] = 24.77, *p* < .001; post-nap: mean = .72, *t*[33] = 18.78, *p* < .001) and shared features (pre-nap: mean = .47, *t*[33] = 10.19, *p* < .001; post-nap: mean = .46, *t*[33] = 10.07, *p* < .001; Fig. 2B). Participants were also above chance in generalization to novel satellites (pre-nap test: mean = .27, *t*[33] = 3.35, *p* = .002; post-nap test: mean = .25, *t*[33] = 2.53, *p* = .017). In both the pre-nap and post-nap tests, performance on unique features was significantly higher than shared features (pre-nap: *t*[33] = 10.24, *p* < .001; post-nap: *t*[33] = 9.32, *p* < .001). This was unexpected given matched performance on these feature types in previous work using a similar paradigm (Schapiro et al., 2017). Analyses of a post-task survey from our study revealed that participants indeed reported focusing on the unique features to solve the task: 61.8% of participants reported using a learning strategy with exclusive focus on unique features, 2.9% reported exclusive focus on shared features, 32.4% reported focus on both unique and shared features, and 2.9% reported no specific focus. Thus, the nature of our specific task setup, which differed from Schapiro et al. (2017) in multiple ways (e.g., satellite names were played out loud instead of displayed on screen), seems to have led participants to focus on the unique features of the satellites during learning.

We assessed whether the focus on unique features during learning was directly associated with worse performance on shared features. For each test and each participant, we calculated the slope of the linear fit that predicted each object’s average shared feature accuracy from its unique feature accuracy. A negative slope would indicate a direct within-object tradeoff between unique and shared feature performance. In the pre-nap test, we did not find evidence of a tradeoff (mean slope = .02, *t*[32] = .37, *p* = .72; Fig. 2C), but this emerged in the post-nap test (mean slope = -.11, *t*[31] = -2.44, *p* = .021; Fig. 2C). A pairwise comparison of pre-nap to post-nap slopes revealed a significant decrease in slope across the nap period (change = -.13, *t*[31] = -2.17, *p* = .038), indicating that a tradeoff in unique and shared feature memory developed over the nap period.

### Real-time TMR cueing drove delayed increases in theta and spindle band power

After completing the pre-nap test, participants were given a 90-min nap opportunity (mean time asleep: 49.26 mins, SD = 18.00 mins; see Extended Data Figure 3-1 for time in each sleep stage). Once they reached N2 sleep, the satellite names were played out loud as TMR cues, with about half of all cues delivered in N2 and half in N3 (mean number of cues presented = 52.82, SD = 28.41; Fig. 3A, Extended Data Figure 3-1).

Previous work demonstrated the importance of administering TMR cue sounds in the up-states of slow-wave oscillations to maximize the likelihood that the cue will trigger a reactivation event (Göldi et al., 2019; Ngo and Staresina, 2022), as the up-state is when endogenous reactivation is thought to occur (Diekelmann and Born, 2010; Mölle and Born, 2011; Gulati et al., 2014; Lewis and Bendor, 2019; Schreiner et al., 2021). In addition, playing a sound immediately after bursts of oscillatory activity known as ‘spindles’ disrupts memory for cued items, perhaps because successful memory reactivation relies on spindle potentiality which is in a refractory state immediately following a spindle (Antony et al., 2018b). To maximize the likelihood of generating reactivation events, we combined these insights to develop a real-time protocol to play TMR cues both in the up-states of slow-oscillations (Ngo and Staresina, 2022) and at least 2.5 seconds after a spindle, outside of the spindle refractory period (Antony et al., 2018b; Fig. 3B, Extended Data Figure 3-2).

The presentation of sound cues led to delayed increases in power in the theta band (cluster 1: 4-7 hz, .60 to 1.20 s after the cue; *p* = .015) and a wide spindle band (cluster 2: 10-21 hz, 1.47 to 2.20 s after the cue; *p* = .007). Both clusters displayed a frontal focus, but neither were lateralized (cluster 1 left vs. right lateralization: *t*[33] = .91, *p* = .37; cluster 2: *t*[33] = .22, *p* = .83; Fig. 3C). These increases in theta and spindle band power replicate work in the TMR literature in which similar responses have been linked to memory reactivation (Schreiner et al., 2015; Cairney et al., 2018; Lewis and Bendor, 2019; Ngo and Staresina, 2022; Liu et al., 2023). We found no differences in neural responses to cues presented in N2 vs. N3 sleep (*p*s > .05). There were no effects of the TMR sound cues beyond the time window displayed in Fig. 3C.

### Cueing drove increases in unique feature memory and decreases in shared feature memory

To assess the impact of reactivation on satellite feature memory while accounting for pre-nap differences in unique and shared feature accuracy and variable numbers of cue repetitions across objects and participants, we applied a mixed-effects model to our behavioral data. The model predicted change in accuracy across the nap for individual items as a function of cueing (cued, uncued), feature type (unique, shared), number of times cued, and pre-nap test accuracy, with participants as random effects factor. We found a strong interaction between cueing and feature type, where cued items exhibited larger increases in unique feature accuracy and decreases in shared feature accuracy than uncued items (β = .17, SE = .045, df = 677.53, *t* = 3.88, *p* < .001; Fig. 4A). Unique and shared feature accuracy change differed for cued items (β = -.19, SE = .03, df = 692, *t* = -5.85, *p* < .001; Fig. 4A) but not for uncued items (β = -.01, SE = .03, df = 683, *t* = -.36, *p* = .72). Cueing significantly increased unique feature accuracy (β (uncued - cued) = -.12, SE = .06, df = 692, *t* = -2.21, *p* = .027; Fig. 4A) and numerically, but non-significantly, decreased shared feature accuracy (β (uncued - cued) = .05, SE = .05, df = 675, *t* = .99, *p* = 0.32). These results were replicated in analyses of confidence ratings (cueing by feature type interaction: β = .26, SE = .12, df = 497.0, *t* = 2.27, *p* = .024).

Novel satellites were all uncued themselves but belonged to (i.e., had overlapping shared features with) categories that were either cued or uncued. Cueing had no direct impact on generalization to novel satellites (β = .04, SE = .04, df = 240.80, *t* = 1.04, *p* = .30).

While the use of mixed-effects models allowed us to control for pre-nap test accuracy of individual items within participants, we wanted to further confirm that the interaction between cueing and feature type was not due to differences in unique and shared feature accuracy before the nap. Thus, we restricted our sample to include only items whose pre-nap shared feature accuracy was greater than or equal to unique feature accuracy (β (shared - unique) = .25, *t*[32] = 6.55, *p* < .001; Fig. 4B). We replicated the significant interaction between cueing and feature type in this sample: cueing predicted increases in unique feature accuracy and decreases in shared feature accuracy compared to uncued items (β = .23, SE = .10, df = 146.51, *t* = 2.28, *p* = .024; Fig. 4B). Unique and shared feature accuracy change differed for cued items (β (shared - unique) = - .23, SE = .08, df = 157, *t* = -2.94, *p* = .004) but not for uncued items (β (shared - unique) = .003, SE = .07, df = 159, *t* = .05, *p* = .96; Fig. 4B). Thus, greater accuracy for unique than shared features in the pre-nap test cannot explain our finding of cueing improving memory for unique features and worsening memory for shared features. We posit that this imbalance in the reactivation benefit may instead be related to either the goals participants formed during learning or the nature of the TMR cues, ideas which we expand on in the Discussion.

Recent work demonstrated that a single TMR cue can impact multiple memories, including memories that are not directly linked to the cue but are highly related to those that are (Schechtman et al., 2021, 2023). Within each cued category we systematically left at least one item out of cueing, resulting in items that were uncued but part of a cued category. We examined whether the impacts of cueing spread to these items by running a model with a cueing term with three levels: uncued items from uncued categories (uncued-uncued), uncued items from cued categories (uncued-cued), and cued items from cued categories (cued-cued). We found a significant interaction between this cueing term and feature type (χ^2^= 15.74, df = 2, *p* < .001; Fig. 4C). For both uncued-cued and cued-cued items, unique feature accuracy was higher and shared feature accuracy lower contrasted to uncued-uncued items (uncued-cued X feature type: β = .11, SE = .05, df = 876.62, *t* = 2.11, *p* = .035; cued-cued X feature type: β = .18, SE = .04, df = 877.28, *t* = 3.96, *p* < .001). Unique and shared feature accuracy change differed for uncued-cued (β (shared - unique) = -.13, SE = .04, df = 883, *t* = -3.04, *p* = .003) and cued-cued (β (shared - unique) = -.19, SE = .03, df = 889, *t* = -6.21, *p* < .001) items, but not for uncued-uncued items (β (shared - unique) = -.02, SE = .03, df = 881, *t* = -.50, *p* = .62). Unique feature accuracy change was significantly greater for cued-cued items compared to uncued-uncued items (cued-cued vs. uncued-uncued: β = -.13, SE = .05, df = 902, *t* = -2.38, *p* = .04; uncued-cued vs. uncued-uncued: β = -.08, SE = .04, df = 891, *t* = -2.13, *p* = .08). Thus, the effects of cueing on unique and shared features spread to uncued items that had cued category-mates.

We explored whether our reactivation findings were impacted by the sleep stage in which cues were delivered (N2 or N3). There was a significant interaction between cueing and feature type, where cueing led to increases in unique feature memory and decreases in shared, for items that were cued only in N2, cued only in N3, and cued in both N2 and N3, when contrasted to items that were uncued (cue time X feature type: χ^2^= 14.96, df = 3, *p* = .002; cued in N2 X feature type: β = .17, SE = .07, df = 665.9, *t* = 2.40, *p* = .017; cued in N3 X feature type: β = .20, SE = .06, df = 665.1, *t* = 3.33, *p* < .001; cued in N2 & N3 X feature type: β = .15, SE = .06, df = 665.6, *t* = 2.55, *p* = .011). Additionally, no significant interaction between sleep stage of cueing and feature type existed when directly contrasting items cued in N2 to items cued in N3 (β = .03, SE = .07, df = 208.0, *t* = .44, *p* = .66). Thus, the sleep stage in which cues were played had no impact on changes in unique and shared feature accuracy. Additionally, we found no relationship between the strength of the neural response to the cues (power in clusters 1 and 2) and the reported changes in unique and shared feature accuracy (*p*s > .08).

### Blocked cue presentation drove greater feature memory shifts than interleaved

To test how the order of reactivation impacts memory, we administered TMR cues in different orders — interleaved vs. blocked — for the two categories of cued satellites (Fig. 3A). We applied the same mixed effects model as above but replaced the cueing (cued, uncued) term with ‘cueing style’ (blocked, interleaved). We found a significant interaction between cueing style and feature type, where blocked cueing led to greater increases in unique feature accuracy and decreases in shared feature accuracy than interleaved (β (interleaved - blocked) = -.11, SE = .06, df = 306.14, *t* = -2.04, *p* = .042; Fig. 5A). Both blocked and interleaved cueing led to differences in unique and shared feature accuracy change (blocked: β (shared - unique) = -.22, SE = .04, df = 316, t = -4.96, *p* < .001; interleaved β (shared - unique) = -.10, SE = .04, df = 315, *t* = -2.36, *p* = .019). Direct pairwise comparisons of blocked vs. interleaved cueing order for shared and unique feature model estimates were non-significant (*p*s > .14). In a model adding uncued items, both blocked and interleaved conditions had a significant interaction with uncued, meaning that cueing differentially impacted unique and shared features even in the interleaved condition (blocked: β = .23, SE = .06, df = 591.35, *t* = 4.05, *p* < .001; interleaved: β = .12, SE = .06, df = 591.23, *t* = 2.11, *p* = .035). We observed no significant differences in sound-elicited time-frequency responses for items in the blocked versus interleaved groups (*p*s > .07).

**Figure 5.**
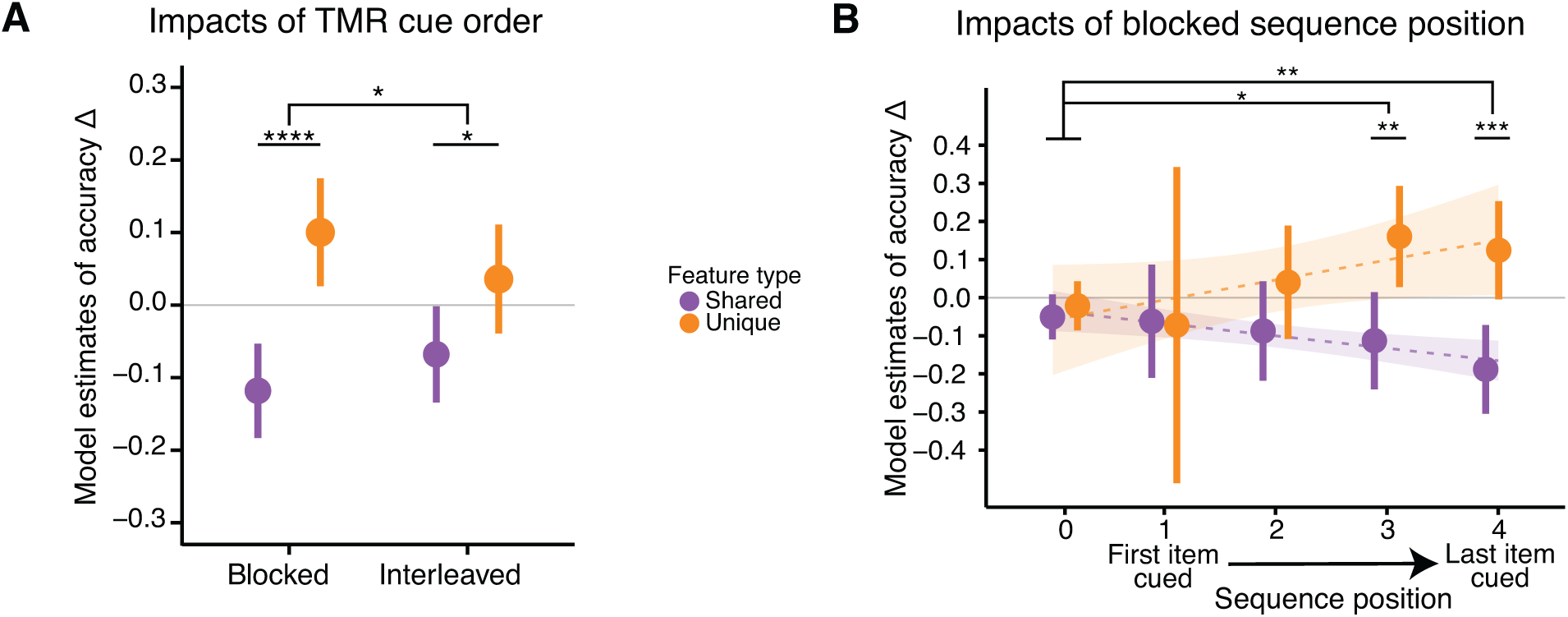
Blocked cue presentation led to greater memory change than interleaved. ***A***, Model estimates of accuracy change for items that were cued in interleaved or blocked order. ***B*,** Model estimates of accuracy change for fully cued blocked items, as a function of their sequence in the cueing order. Sequence position 4 corresponds to the last item cued before the participant woke up from their nap; 0 corresponds to items in the uncued category. Plotted are model estimates for accuracy change for unique and shared features, with a linear fit to sequence position (shaded area = 95% confidence intervals).

We additionally investigated whether blocked reactivation impacted all items equally across sequence position (first, second, third, or fourth cued position). We replaced the cueing (cued, uncued) term from our model with ‘sequence position’ (0, 1, 2, 3, 4) and ran this model on fully cued blocked items. A sequence position of 4 represented the last item cued before the end of the nap; 3 represented the second to last item cued; 2 represented the third to last item cued; 1 represented the fourth to last item cued; 0 represented items from the uncued category. We found a significant interaction between sequence position and feature type (β = .07, SE = .02, df = 361.59, *t* = 3.66, *p* < .001), where shared feature accuracy decreased across sequence positions (trend = - 0.03, SE = .01, df = 362, lower CL = -.056, upper CL = -.003) and unique feature accuracy increased across sequence positions (trend = .04, SE = .01, df = 369, lower CL = .01, upper CL = .07; Fig. 5B). We next converted sequence position to a categorical term in the model to investigate the individual sequence position levels. We again report a significant interaction between sequence position and feature type (χ^2^= 13.65, df = 4, *p* = .009) whereby blocked sequence positions 3 and 4 drove increases in unique accuracy and decreases in shared accuracy contrasted against uncued items (sequence position 3: β = .24, SE = .10, df = 353.92, *t* = 2.50, *p* = .013; sequence position 4: β = .28, SE = .09, df = 356.09, *t* = 3.04, *p* = .003; Fig. 5B). There was no significant interaction between sequence position 1 or 2 and feature type (shared, unique) contrasted against uncued items (*p*s > .35). Unique and shared feature accuracy change differed for blocked sequence position 3 (β (shared - unique) = -.27, SE = .09, df = 357, *t* = -2.97, *p* = .003) and 4 (β (shared - unique) = -.31, SE = .09, df = 360, *t* = -3.59, *p* < .001), but not for positions 1, 2, or uncued items (*p*s > .20). Neural responses were similar across blocked sequence positions (*p*s > .25), suggesting that the effect of sequence position was not due to differential cue processing across the nap.

## DISCUSSION

Our memories are not exact recapitulations of our original encoded experiences; rather, they transform across time, with certain features of our memories becoming stronger while others fade away. We demonstrate a role for memory reactivation during sleep in this transformation process. We found that reactivating individual novel satellite objects during sleep differentially impacted memory based on feature type: cueing improved memory for features unique to individual satellites and worsened memory for features shared across satellites in the same category. Thus, offline memory reactivation does not always act holistically on object memories. These results echo findings that memories for multi-featural objects are not represented as bound units, as different features can be independently remembered or forgotten (Brady et al., 2013; c.f., Balaban et al., 2020). Our work suggests that reactivation during sleep may be a mechanism supporting such selective, independent feature remembering and forgetting.

We found evidence that reactivation produced a competitive dynamic between unique and shared feature memory, with memory for unique and shared features within the same object becoming negatively correlated across the nap. The overall improvement of unique features and deterioration of shared features, in addition to evidence of a direct trade-off, are suggestive of an underlying differentiation process where enhanced distinguishing features and suppressed overlapping features reflect the separation of exemplar representations (Fig. 6). Such a differentiation process would explain our finding that cueing effects spread to uncued category members: movement of cued exemplars away from the central tendency of the category would diminish that central tendency for uncued exemplars and reduce interference for the uncued item’s unique features. These reactivation-driven, competitive dynamics are reminiscent of those discussed in the Retrieval Induced Forgetting (RIF) literature, wherein retrieval of certain associations drives the forgetting of related associations, perhaps in the service of differentiation (Anderson et al., 1994; Anderson and Spellman, 1995; Norman et al., 2007). Awake reactivation has been proposed to play a role in RIF effects (Wimber et al., 2015) and orthogonalization (Nguyen et al., 2023), and the connection between RIF and sleep reactivation was investigated in a recent TMR study (Joensen et al., 2022): participants studied overlapping pairs of items, where one pair was encoded more strongly than the other. Joensen et al. (2022) found that TMR during a subsequent nap enhanced memory for the strongly-encoded pair and degraded memory for the weakly-encoded pair (see also Oyarzún et al., 2017). Our work shows that competitive dynamics can play out between different kinds of features within an individual item. In our case, these dynamics are not explained by differences in encoding strength. Thus, there may be multiple factors that interact to determine which memories are enhanced versus suppressed following sleep reactivation.

**Figure 6.**
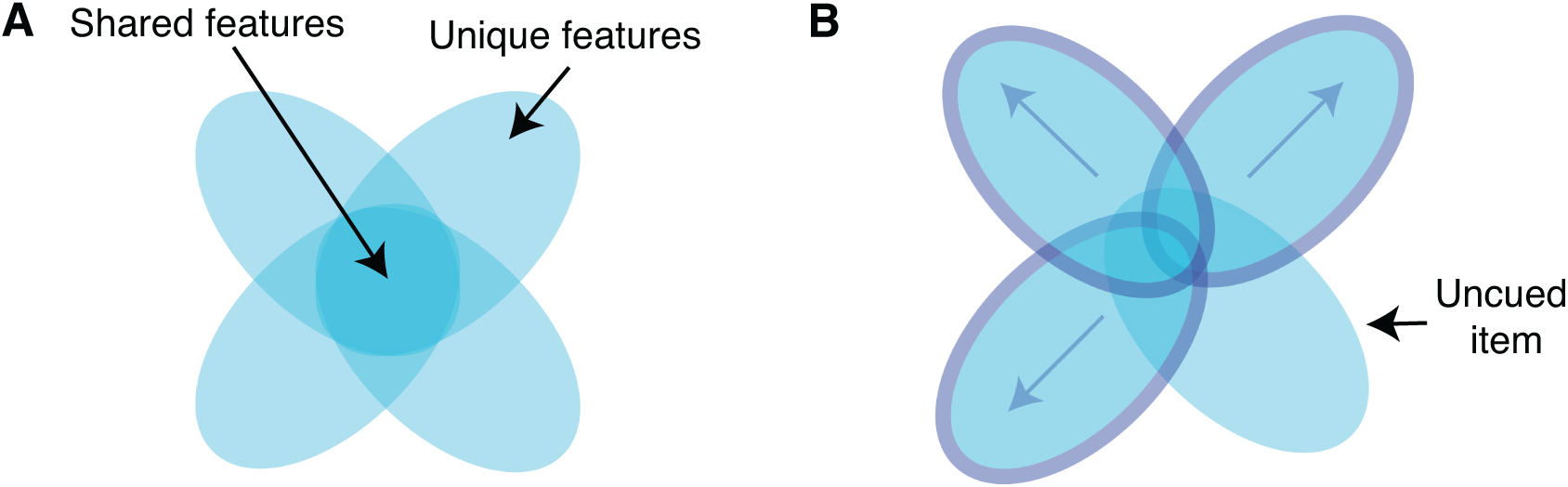
Cueing as a driver of differentiation. ***A*,** Schematic of representational overlap for four exemplars from the same category. ***B*,** If cued objects differentiate, the unique aspects of their representations are enhanced and the shared aspects are diminished. These effects generalize to an uncued item, because the differentiation of the other exemplars reduces overlap for all items, and all unique elements are subject to less interference.

Across our analyses, we found that sleep reactivation benefitted unique features. The Complementary Learning Systems (CLS) theory proposed that offline hippocampal reactivation enables the extraction of shared structure across memories in neocortex (McClelland et al., 1995). In support of this theory, sleep has been shown to benefit structure memory, the abstraction of gist information, memory integration, and insight (Wagner et al., 2004; Ellenbogen et al., 2007; Durrant et al., 2011; Lewis and Durrant, 2011; Landmann et al., 2014; Schapiro et al., 2017; Lerner and Gluck, 2019). In contrast, other work has suggested that sleep is critical for pattern separation (Mirković and Gaskell, 2016; Saletin et al., 2016; Hanert et al., 2017; Doxey et al., 2018), memory specificity (Barry et al., 2019), and the protection of individual memories from interference (Ellenbogen et al., 2006; McDevitt et al., 2015). There have also been failures to find a relationship between sleep and memory generalization (Mirković and Gaskell, 2016; Hennies et al., 2017; Schapiro et al., 2017; Tandoc et al., 2021). One potential way of reconciling this discrepancy in the benefits of sleep for general versus specific information is that reactivation may impact learning locally within the hippocampus in addition to potential impacts in the neocortex. Our finding of sleep reactivation benefitting unique features could potentially reflect this local hippocampal learning. Additionally, REM sleep may sometimes play a role in observed benefits to memory structure and integration (Cai et al., 2009; Sterpenich et al., 2014; Tamminen et al., 2017; Lewis et al., 2018). Future work is needed to disentangle the factors that lead to different kinds of memory changes and learning in different memory systems across studies, but below we offer two explanations for patterns observed in our particular case.

The consolidation trajectory hypothesis posits that sleep serves to reinforce goals formed during learning, such that if interconnected elements of a complex memory are learned harmoniously, they will be jointly promoted during sleep; if they are learned antagonistically, they will be pitted against each other during sleep (Antony and Schechtman, 2023). One prior TMR study demonstrated that generating specific competitive dynamics during pre-nap learning modulated the impacts of reactivation (Antony et al., 2018a). In our study, reactivation-driven competitive dynamics may have favored unique features because participants prioritized learning of unique features over shared features before the nap. However, while participants reported focusing on, and exhibited better memory for, unique than shared features pre-nap, we did not observe a within-object tradeoff in unique and shared feature accuracy pre-nap. Antony & Schechtman (2023) highlight, though, that it is unknown whether competitive dynamics need to be explicitly present before sleep or whether reactivation can drive these dynamics itself. Our findings lend support to the idea that reactivation can drive competitive dynamics between features of object memories based on pre-sleep learning goals, even if the features are not yet in direct competition.

An alternative interpretation of our findings is that reactivation drove selective improvements to unique features because the reactivation cues were themselves unique features (the individual satellite names). Indeed, a prior TMR study using category labels as cues found that cueing led to worse detail memory but not worse category memory (Witkowski et al., 2021). Thus, the way information is processed during reactivation may depend on the match between the reactivation cue and memory trace (akin to synergistic ecphory accounts of retrieval; Tulving, 1982). In our study, the use of a unique feature cue may have led to stronger reactivation of unique than shared information, benefitting the individuating features of the objects at the expense of the shared features. In summary, while future work is needed to resolve the factors that modulate whether reactivation benefits unique or shared features, our results provide a proof of concept that reactivation can modulate individual features of objects differentially and could thus be driving sleep-dependent memory transformation.

In addition to assessing how reactivation impacts different features of an object, we investigated whether the order of this reactivation matters. While both blocked and interleaved reactivation orders led to improvements in unique and decrements to shared feature memory, the effects were especially strong for items reactivated in blocked order. Our findings align with recent work demonstrating that blocking can drive differentiation of memory representations (Flesch et al., 2018, 2023). Interestingly, Flesch et al. (2018) found that blocking was most beneficial to participants who already had an existing internal model that emphasized the dimensions on which stimuli were later differentiated, and the impacts of blocking could be replicated in neural networks if they were given a differentiated model to build upon. Thus, perhaps a combination of blocked reactivation with a pre-nap internal model emphasizing unique features led to the strong differentiation of object memories within categories in our study. Note, though, that this work, as in other work from the category learning literature, manipulated the order of categories, whereas we manipulated the order of items within a category. But the correspondence across literatures points to a general benefit of blocking for differentiation, which could be exploited in future studies aiming to improve memory for confusable information.

We found the strongest memory effects for items cued later in the blocked series. This could be due to either retroactive interference, such that memory improvements of early-cued objects were dampened by subsequent reactivations of similar objects, or simple decay over time. This examination of cueing order was inspired by CLS, which proposed that interleaved memory reactivation should benefit shared features (in the context of neocortical learning, as discussed above) and that blocking could lead to interference, as repeated reactivation of an item could overwrite representations of similar items presented earlier (McClelland et al., 1995). Though we did not find overall reactivation benefits for shared features, we did find interleaving to be less detrimental to shared features than blocking. Additionally, retroactive interference during blocking would align with CLS principles. More work is needed to understand the potential interactions between reactivation order, memory types, and task characteristics that lead to these patterns of behavior.

Our work provides a potential explanation for how object memories transform over time. Recent work suggests that memory for episodes may not undergo the same kind of transformation: Andermane et al. (2023) reported that while the features of an object were forgotten independently of one another, if an event was composed of three unrelated items (i.e., an object, location, and person), the items making up an event were remembered or forgotten jointly (Horner et al., 2015; Joensen et al., 2020; Andermane et al., 2023). However, other work has demonstrated that the unique features of an episode can be forgotten while the gist is retained (Winocur and Moscovitch, 2011; Sekeres et al., 2016, 2018b). Thus, the types of features comprising an event and how they relate to one another may impact whether they are remembered jointly or independently, and the extent to which the event is transformed. While our work highlights that reactivation has the power to act on single entities non-holistically, future work is needed to establish under what conditions reactivation may drive analogous transformations to events as seen here with objects.

We developed a novel protocol for this study to play sound cues in moments optimized to generate reactivation events: both in the up-states of slow oscillations (e.g., Ngo & Staresina, 2020) and outside of the spindle refractory period (Antony et al., 2018b). Using this protocol, we replicate previous findings showing that TMR cue delivery triggers delayed increases in theta and spindle band power (Cairney et al., 2018; Göldi et al., 2019; Ngo and Staresina, 2022; Wick and Rasch, 2023), with the delayed spindle increase coinciding with the following upstate (Ngo and Staresina, 2022). This modulation in spindle power may correspond to item reactivation that is evoked in the upstate following the cued one (Belal et al., 2018; Cairney et al., 2018; Ngo and Staresina, 2022; Abdellahi et al., 2023). We found similar neural and behavioral responses to cues presented across N2 and N3 sleep, replicating reports that both N2 and N3 cueing lead to memory benefits (Hu et al., 2020). This lack of difference across N2 and N3 cueing in our study is likely due to targeting up-states regardless of sleep stage. If cues were instead delivered randomly, they may be presented in different neural states and generate different responses across N2 and N3 sleep (Carbone et al., 2023; Wick and Rasch, 2023). In other words, specific sleep features like slow oscillations and spindles may be key to understanding memory reactivation, beyond and regardless of the sleep stage they occur in.

In sum, our findings suggest that memory reactivation during sleep can differentially impact features within an object — reactivation does not always act holistically on object memory. Cueing individual objects enhanced memory for distinguishing features at the expense of features shared with other members of an object’s category. We interpret these non-holistic, competitive dynamics to reflect a differentiation of item representations driven by sleep-based reactivation. While future work will be needed to determine under what stimulus (e.g. events vs. objects) and task conditions (e.g., encoding focus on distinctive vs. shared features) these effects will manifest, our results provide a proof of concept that reactivation in the sleeping brain need not be holistic. Our findings emphasize the multidimensional nature of our memories and the changes they undergo over time and suggest that memory reactivation during sleep may be a critical mechanism driving memory transformation.

## ACKNOWLEDGEMENTS

The authors thank Sirisha Krishnamurthy for help with data collection, Dr. Cybelle Smith for assistance setting up the lab EEG system and for EEG analysis discussions, and Dr. Hong-Viet V. Ngo for sharing code for real-time sound delivery. This work was supported by NSF GRFP DGE-2236662 to E.M.S., NIH grant R01 MH129436 to A.C.S, and Charles E. Kaufman Foundation grant KA2020-114800 to A.C.S.

## EXTENDED DATA

**Figure 3-1.**
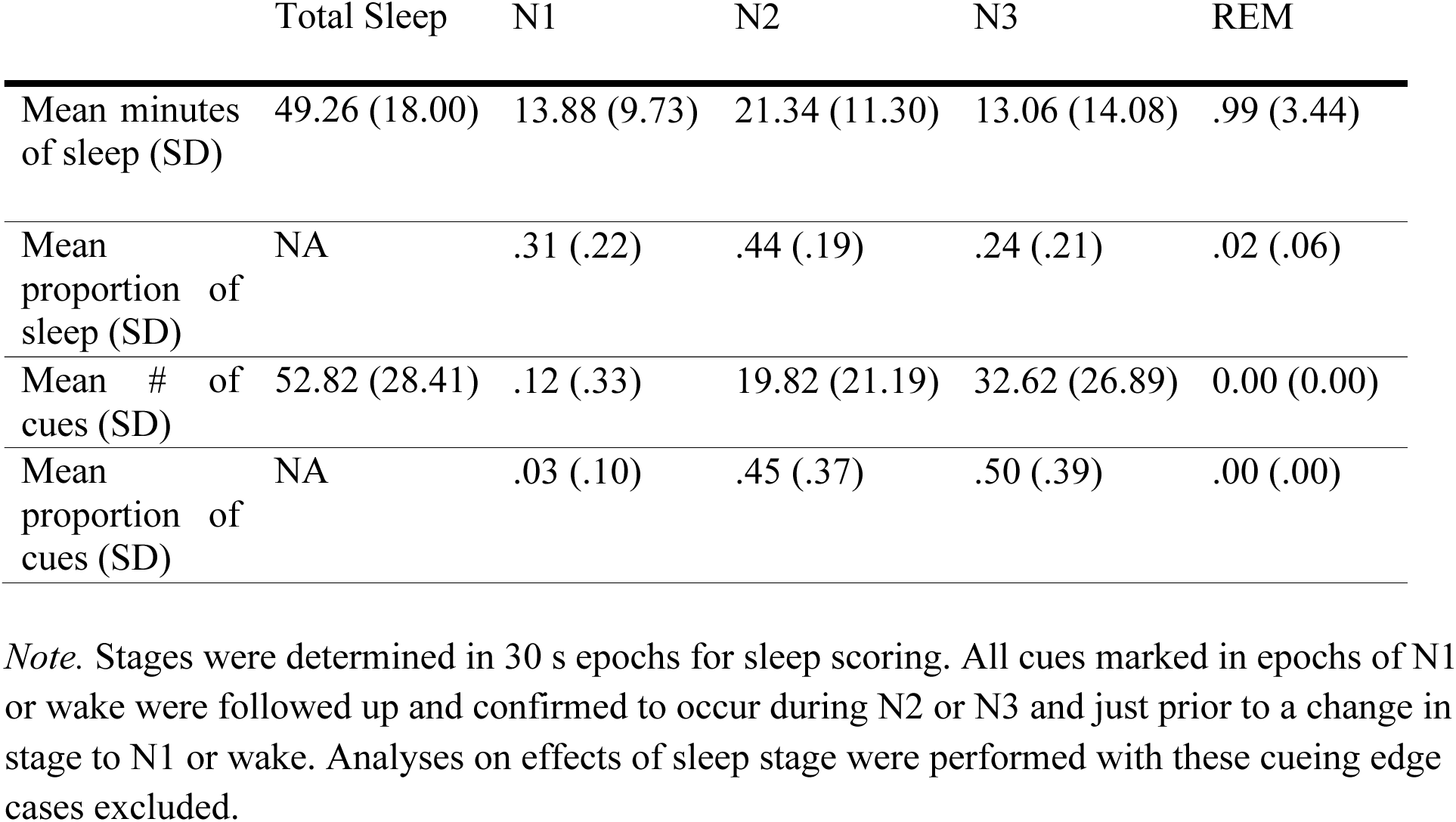
Table of average sleep time and cue numbers.

**Figure 3-2.**
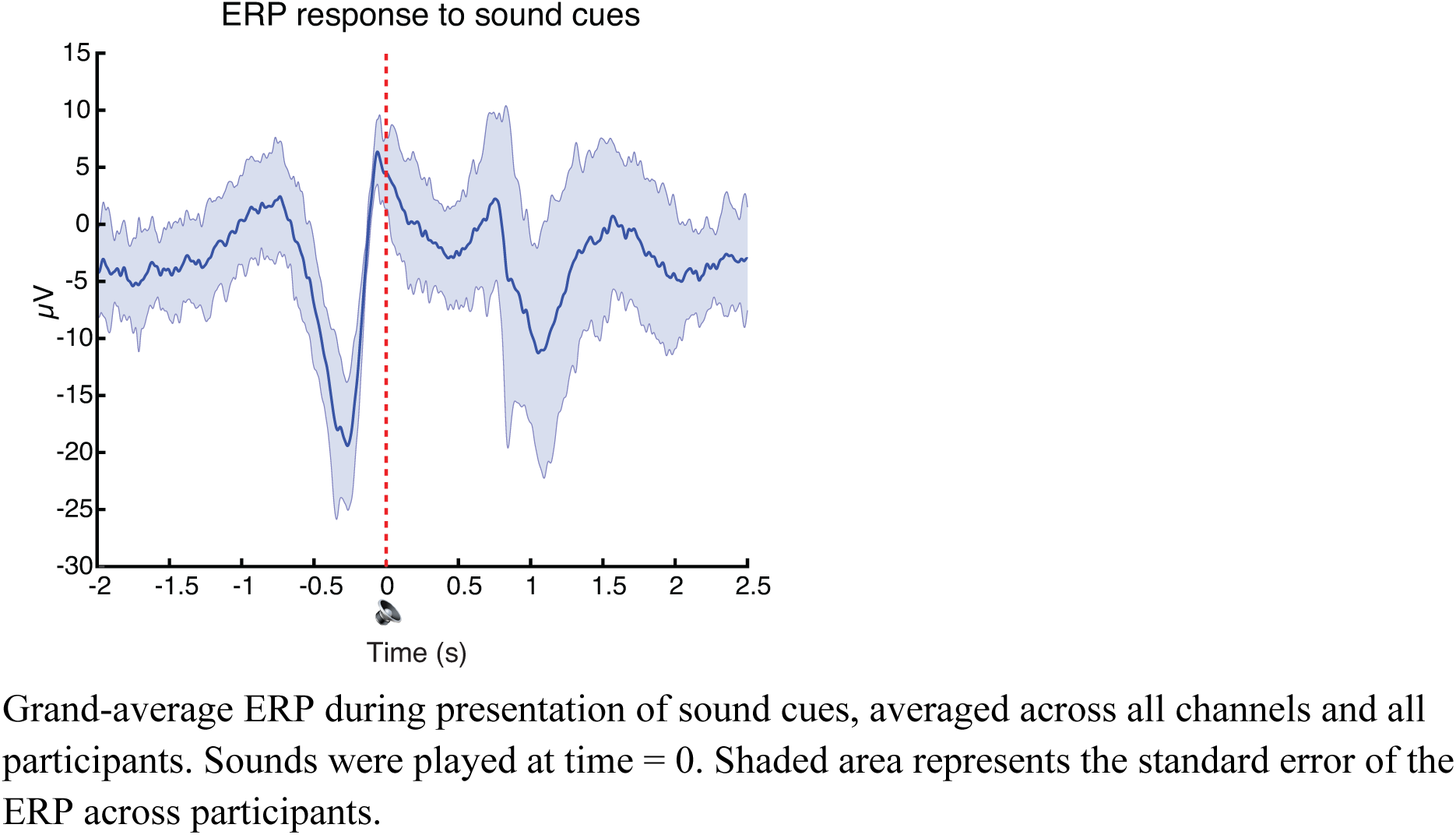
Sounds were consistently delivered in SO up-states.

**Figure 4-1.**
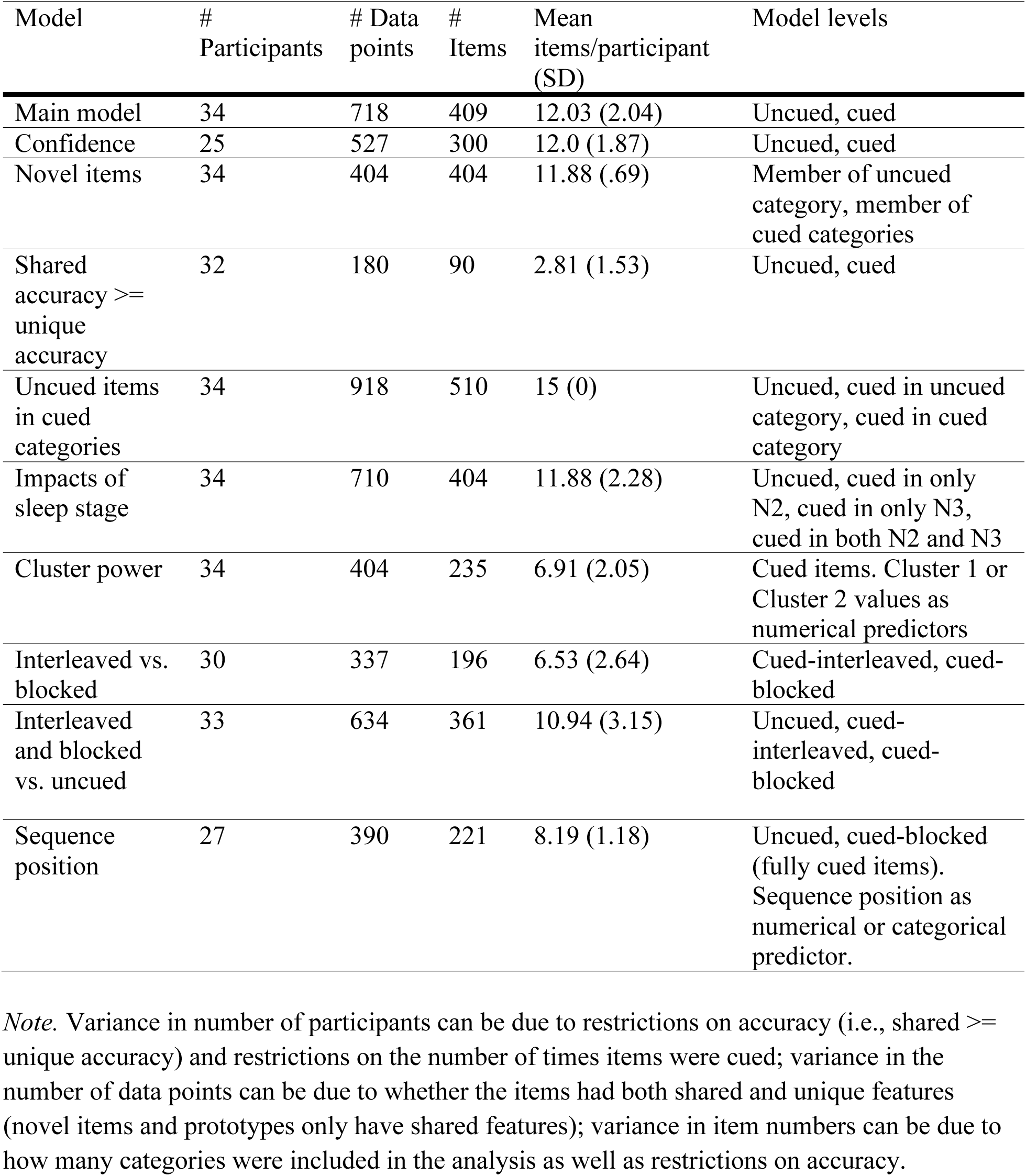
Table of model details.

